# Establishment of an antimetabolite-based transformation system for the wood-decaying basidiomycete *Phanerochaete chrysosporium*

**DOI:** 10.1101/2025.06.05.658208

**Authors:** Kazuma Masumoto, Petra Banko, Ayane Yamamoto, Kyoko Miwa, Chiaki Hori

## Abstract

The model wood-decaying basidiomycete Phanerochaete chrysosporium has been extensively studied to elucidate the molecular mechanisms of wood decomposition. However, genetic studies have been limited by the lack of adequate genetic tools. Here, we established an antimetabolite-based transformation system, originally developed for ascomycetes, for use in P. chrysosporium. The transformation system utilizes pyrithiamine (PT), a thiamine antimetabolite, in combination with the pPTRII vector that contains the PT resistance gene (ptrA). PT effectively inhibited the growth of P. chrysosporium, and the introduction of ptrA conferred resistance to transformant mycelia. The transformation efficiency was comparable to that in ascomycetes, suggesting that the transformation system is also applicable to basidiomycetes. To examine the suitability of the system for heterologous gene expression, four cassettes were constructed to express GFP under the promoters of the actin1, DED, and GAPDH genes. Promoter activities were assessed via fluorescent microscopy observation of transformant mycelia and GFP quantification in crude cell extracts, revealing that the actin1 promoter drove the highest level of expression. Furthermore, truncating repeat sequences of the autonomously replicating sequence in the vector backbone improved transformation efficiency, likely due to the reduction in vector size. The transformation efficiency of the gene cassette-inserted vector in P. chrysosporium was relatively higher than that reported with alternative transformation systems in other species of wood-decaying basidiomycetes. The present transformation system could provide a platform for protein expression and genetic engineering in P. chrysosporium and potentially in other wood-decaying basidiomycetes.

**Importance:** Wood-decaying basidiomycetes are well-recognized for their exceptional capabilities to decompose lignocellulosic biomass and oxidize a broad range of complex organic compounds. These capabilities are essential for maintaining the forest ecosystem and hold potential in biotechnological applications such as transforming recalcitrant biomass into useful compounds and degrading toxic substances in industrial effluents. However, genetic manipulation in basidiomycetes remains challenging because of the inefficiency of transformation systems. In the model lignocellulose-degrading basidiomycete, *P. chrysosporium,* transformation methods using dominant markers are scarce and were reported over two decades ago, necessitating the re-establishment of a functional system compatible with modern genetic tools. In this study, an efficient genetic transformation system was achieved by using an antimetabolite-based selection strategy for *P. chrysosporium.* This transformation system would lay the foundation for advancing our understanding of the molecular mechanisms of wood decomposition and support the targeted optimization of basidiomycetes for various biotechnological applications.

## INTRODUCTION

Wood-decaying basidiomycetes are well-recognized for their exceptional capabilities to decompose recalcitrant biomass and oxidize a broad range of complex organic compounds (1, 2). *Phanerochaete chrysosporium* has been widely used as a model fungus of white-rot due to its high lignocellulose degradation capacity, rapid growth rate, and sporulation ability (3–5). It efficiently decomposes plant cell wall components such as cellulose, hemicellulose, and lignin by secreting various carbohydrate-active enzymes (CAZymes) (6). These lignocellulose-decomposing enzyme systems have been well studied through biochemical characterization and genome-wide expression analyses, including transcriptomics and proteomics (7). Moreover, several comparative genomic studies of wood decay fungi, including *P. chrysosporium*, also provide insights into their diversity and evolution in lignocellulose degradation (2, 8, 9). However, in basidiomycetes, genetic tools, including overexpression systems and genome editing, are comparatively underdeveloped.

Two main selection strategies have been commonly utilized for transforming *P. chrysosporium*. The auxotrophic selection method was developed following the isolation of uracil and adenine auxotrophic mutant strains in the 1980s (10–13). However, the generation of these mutant strains is labor-intensive and can negatively impact growth (14). Other methods utilize dominant selectable marker genes that confer resistance to antibiotics, such as G418 (Geneticin) (15) and phleomycin (16), or herbicides like bialaphos (14). This approach enables the transformation of wild-type strains while also circumventing the need for strain development (14). Nonetheless, these transformation methods were reported more than two decades ago. Therefore, developing an optimized transformation pipeline in *P. chrysosporium* is essential to progress genetic research and translational applications.

In ascomycetes, the pyrithiamine (PT) resistance gene, *ptrA*, was used as a dominant selectable marker, establishing an antimetabolite-based transformation method as an alternative for conventional antibiotic selection (16). PT acts by replacing thiamine (vitamin B12) in cells, causing thiamine deficiency and generating toxic byproducts that interfere with energy-producing pathways such as glycolysis and the citric acid cycle (17). The *ptrA* gene, derived from *Aspergillus oryzae,* encodes a predicted thiamine synthase(18), which likely confers PT resistance through the overproduction of thiamine (18). Several *ptrA*-based vector systems emerged in recent years, broadening the genetic resources available for genome editing in ascomycetes (19–21). However, to date, there are no reports of using PT-selection in basidiomycetes.

In this study, a novel transformation system for *P. chrysosporium* was established using the antimetabolite PT and a plasmid carrying the *ptrA* resistance gene. A series of expression vectors harboring the *egfp gene* under various native promoters was constructed to select a suitable promoter for heterologous expression. Finally, the transformation efficiency was enhanced by deleting one of the repeat sequences of the autonomous replication sequence present in the plasmid. Our work should provide a foundation for further development of efficient transformation systems of wood-decaying basidiomycetes and functional genetic studies to help address knowledge gaps in the metabolic and regulatory networks of *P. chrysosporium*.

## RESULTS

### PT inhibits the growth of *P. chrysosporium*

First, to verify whether PT inhibits the growth of the wild-type *P. chrysosporium*, protoplasts were plated on CD agar plates supplemented with PT concentrations ranging from 0.1 to 1.0 µg/mL. A decrease in the number of recovered colonies, proportional to the increase in PT concentration, was observed (Supplementary Table S1). Next, protoplasts were transformed via the PEG method with 1 µg pPTRII and cultured on CD agar plates supplemented with 0.3 µg/mL PT. After the transformation procedure, greater mycelial growth was observed, compared to the plate with the wild-type strain grown under the same conditions (Fig. 1A), suggesting that the *ptrA* resistance gene is a selectable marker gene in *P. chrysosporium*.

**Figure 1.**
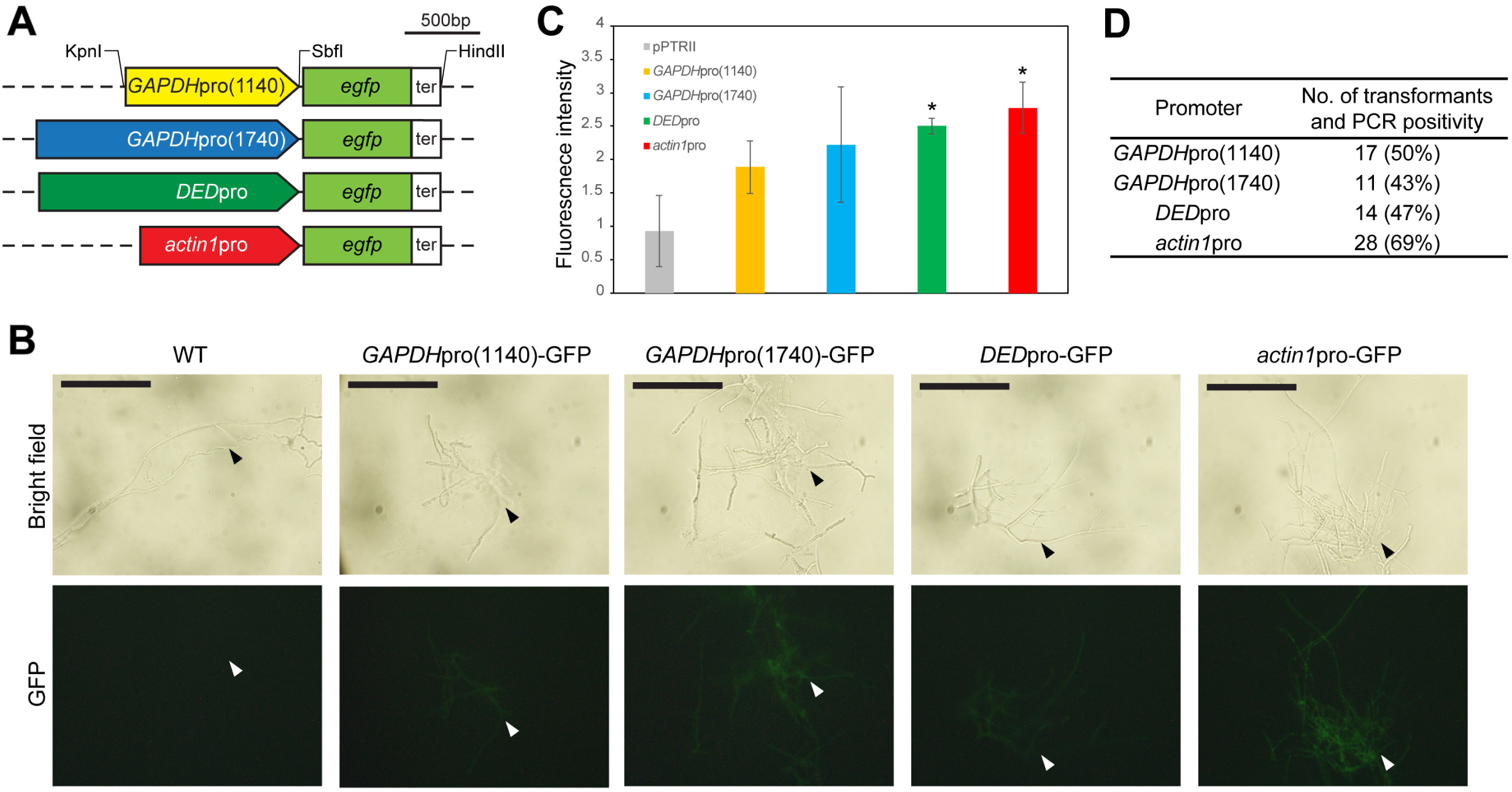
Establishment of the PT selection method for *P. chrysosporium* transformation using the pPTRII vector. **A)** Agar plugs of the wild-type (WT) and pPTRII transformed mycelia were cultured for 4 days on CD agar plates containing 0.3 µg/ml PT. WT showed susceptibility to PT, but the transformants grew and exhibited PT resistance. **B)** Genomic PCR using primers targeting the pPTRII backbone was performed to confirm transformation in PT-resistant colonies (1–8). The ITS region was amplified as a control for PCR conditions. **C)** Effects of vector amount and PT concentration in the selection media on the number of transformants and PCR positivity. The number of transformants was estimated from the total colony count per 10^7^ protoplasts, adjusted based on the proportion of PCR-confirmed positives among seven randomly selected colonies.

To verify the presence of the pPTRII plasmid in PT-resistant colonies, genomic DNA isolates from mycelia were subjected to PCR using vector-specific primers. The internal transcribed spacer (ITS) region was amplified as a reference control for the genomic DNA amount. Of the eight randomly selected colonies, three produced the expected amplicon (Fig. 1B). This result indicated that at least multiple independent cells were transformed with pPTRII, resulting in PT resistance. However, a high rate of false-positive colonies was observed, as many PT-resistant colonies lacked the vector.

To reduce the rate of false positives, the optimal transformation conditions were determined with PT concentrations ranging from 0.1 to 1.0 µg/mL, and donor DNA concentrations of 1, 5, and 10 µg. The number of transformants was estimated from the total number of PT-resistant colonies and the PCR positivity rate (Fig. 1C). A high frequency of PCR negatives was observed at a donor DNA concentration of 1 µg, regardless of the PT concentration. Increasing the donor DNA amount to 10 μg, significantly improved the number of transformants, accompanied by a decrease in the number of false positives. The highest number of transformants was obtained at 0.3 μg/mL PT, where transformation of 1 × 10^^7^ protoplasts with 10 µg of donor DNA yielded 96 transformants, with PCR confirming the presence of plasmid in all seven tested colonies. These conditions were therefore utilized for all subsequent transformation experiments.

### Native promoters drove GFP expression in transformants

With the established transformation protocol, we aimed to design and optimize a heterologous gene expression vector by incorporating a GFP reporter cassette in the pPTRII vector. Four GFP expression cassettes were constructed from the native promoter regions of three housekeeping genes: *actin1, GAPDH*, and *DED* (Fig. 2A). *Actin1* is a housekeeping gene present in all eukaryotes (22, 23) and has also been used as an internal control for expression in *P. chrysosporium* (24). The *GAPDH* promoter (1,140 bp) has been previously evaluated in *P. chrysosporium* for reporter gene expression but exhibited fluorescence at only background levels (25). We speculated that including the TATA box might improve promoter activity, therefore, we extended the promoter 1,740 bp upstream of the *GAPDH* start codon. In the mushroom-forming basidiomycete, *Coprinopsis cinerea,* the *DED* promoter has been reported to drive higher promoter activity than other constitutive promoters (26). Therefore, we identified a homolog of the *DED* gene in the *P. chrysosporium* genome and selected its upstream region as a promoter candidate. Each promoter region was amplified from genomic DNA of *P. chrysosporium* and fused upstream of the *egfp* gene, followed by the terminator region of the *GAPDH* gene. The assembled cassettes were inserted into pPTRII in the KpnI and HindIII restriction sites (Fig. 2A).

**Figure 2.**
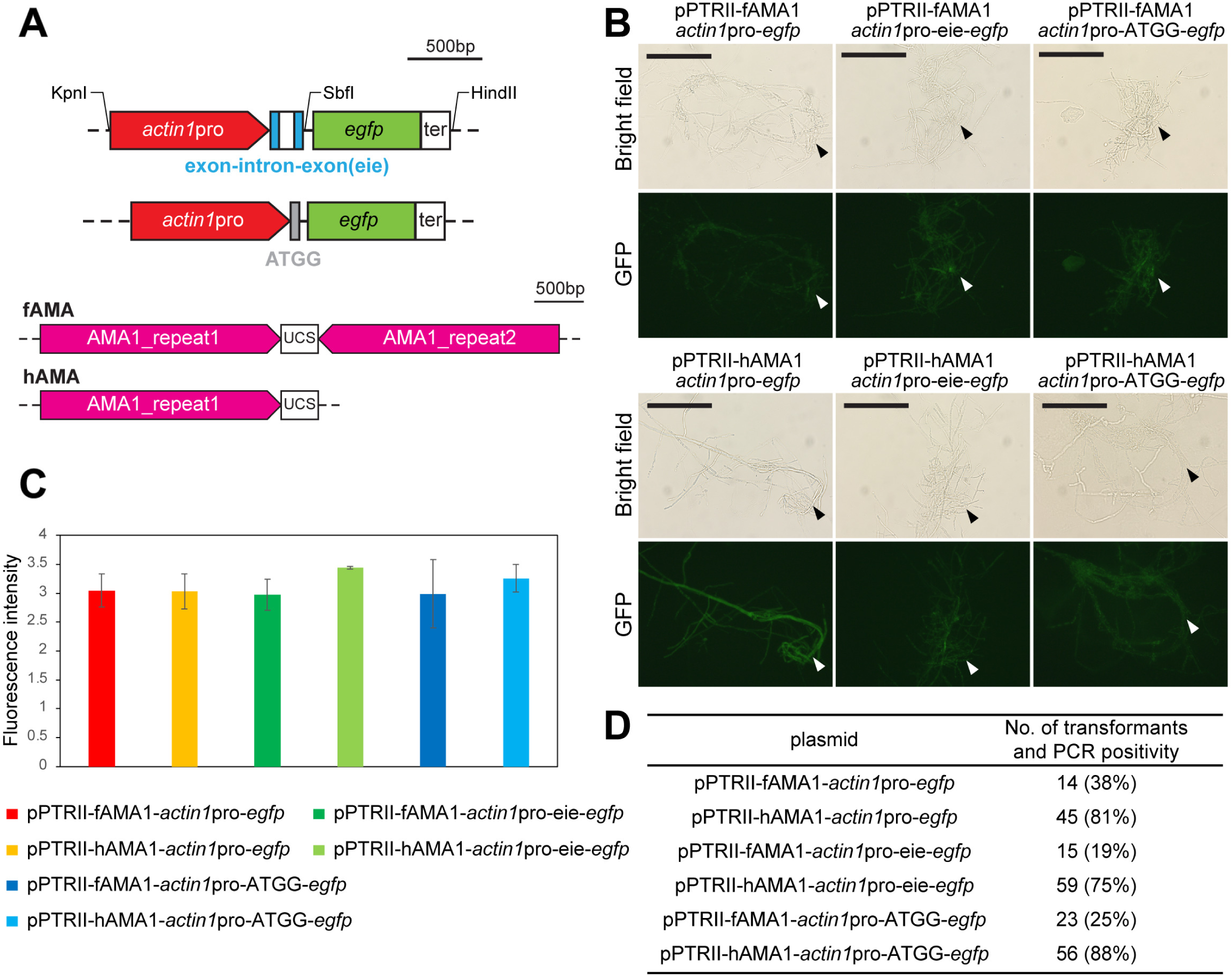
Evaluation of promoter activity via GFP reporter expression in *P. chrysosporium.* **A)** Schematic representation of the four GFP expression cassettes, composed of the respective homologous promoter regions, the GFP coding region, and the *P. chrysosporium GAPDH* terminator region (ter). Restriction sites utilized during the cassette construction and cloning are indicated. Sequence lengths are proportional to the scalebar. **B**) Fluorescent microscopy of mycelia transformed with GFP vector constructs. The top panel shows the bright field, and the bottom panel shows the fluorescent micrographs of transformed mycelia. Scale bar: 100 µm. **C)** Comparison of GFP fluorescence intensity in the crude intracellular protein of GFP transformant mycelia driven by different promoter regions. Mean ± SD values from three biological replicates are presented. Statistical difference was calculated by Welch’s t-test (*, p<0.05). **D)** The number of transformants with each GFP vector construct. The number of transformants was estimated from the total colony count per 10^7^ protoplasts, adjusted based on the proportion of PCR-confirmed positives among 16 randomly selected colonies. Transformations were performed with 10 µg of each GFP expression vector, and protoplasts were cultured on CD agar media containing 0.3 µg/ml PT.

*P. chrysosporium* protoplasts were transformed with the GFP expression vector constructs. To verify the expression of the introduced GFP cassettes, the hyphae of the transformants were observed using fluorescence microscopy (Fig. 2B). GFP signal was detected in the hyphae of all four transformants, while the wild-type strain showed no detectable fluorescent signal.

To assess promoter activity, GFP expression levels were quantified in 1 mg/mL crude intracellular proteins using fluorescence spectroscopy. The mycelia transformed with vectors containing the *DED* and the *actin1* promoters showed significantly higher intracellular GFP fluorescence intensity compared to the empty vector control (Fig. 2C). Although the extended *GAPDH* promoter showed variation in the promoter activity, the fluorescence signal was not statistically different compared to the reported original *GAPDH* promoter. The *actin1* promoter showed the highest mean fluorescent intensity levels among the four tested promoters, approximately 1.5 times more than *GAPDH*pro(1140), which exhibited the weakest promoter activity. Similarly, the vector containing the *actin1* promoter yielded the highest number of transformants (Fig. 2D) and the highest PCR positivity rate calculated from 16 randomly selected colonies (Supplementary Fig. S1). Therefore, we selected the *actin1* promoter construct for further optimization steps.

### Shortened AMA1 sequence improved transformation efficiency

Additional modifications were introduced to both the cassette and the vector backbone of the pPTRII-*actin1*pro-*egfp* construct (Fig. 3A). Improved gene expression levels have been previously achieved by inserting the conserved Kozak sequence in filamentous fungi (27) or introns in the gene coding sequences in *P. chrysosporium* (25). Here, two sequences, an exon-intron-exon sequence derived from the *GAPDH* gene and a four-nucleotide ATGG sequence, were examined (Fig. 3A, top). The pPTRII vector contains AMA1, an autonomously replicating sequence commonly used in filamentous fungi (28, 29). It consists of two inverted repeat sequences connected by a unique central spacer sequence (UCS), spanning over 5.2 kb (29, 30). Previous reports suggested that shortening this region is beneficial for handling and vector construction, while having relatively low impact on transformation efficiency (31). Therefore, a half AMA1 region (hAMA) containing a single repeat and the UCS region was constructed by inverse PCR and compared with the full-length version of AMA1(fAMA) (Fig. 3A, bottom).

**Figure 3.**
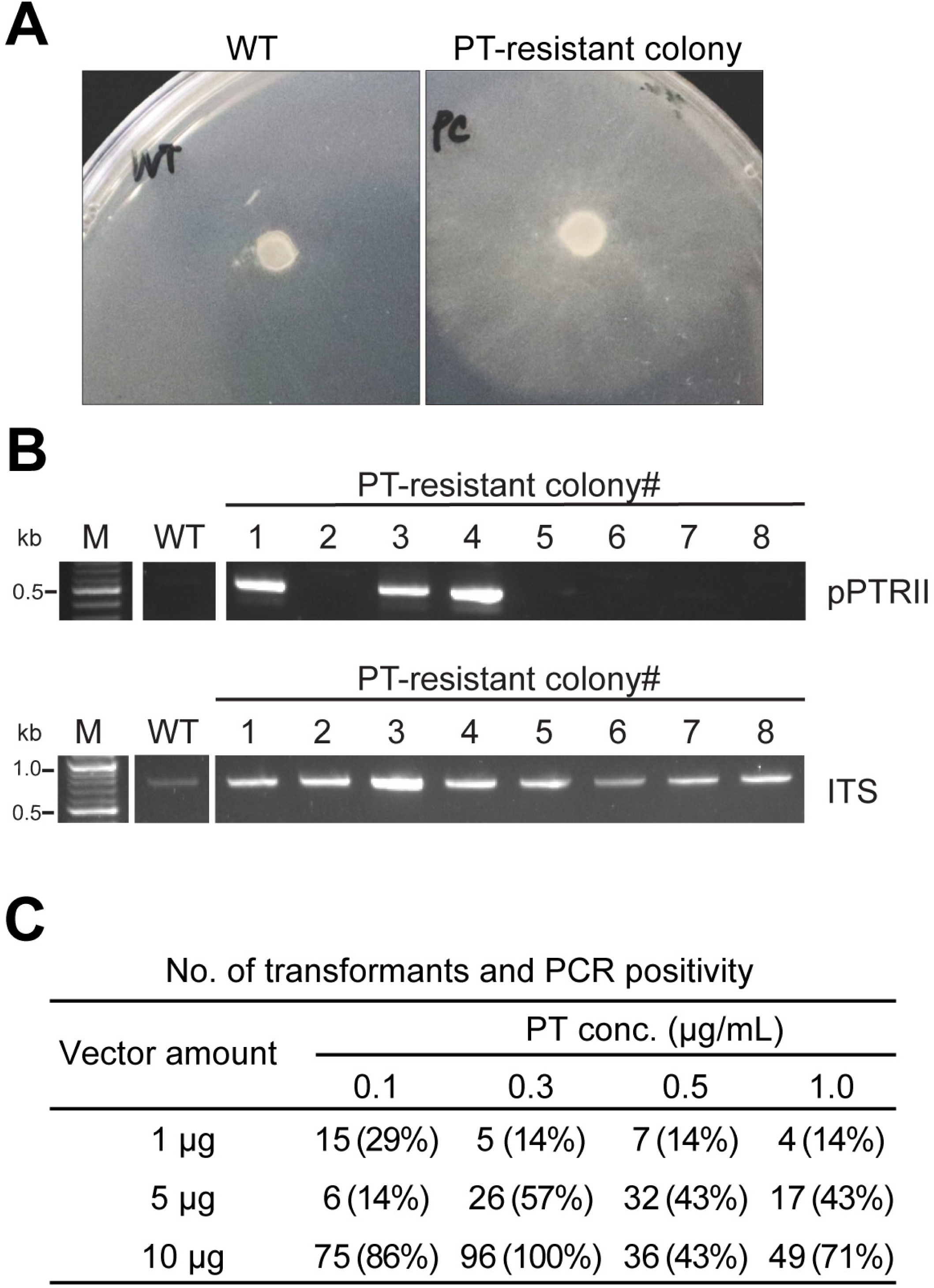
Modified vector constructs for improving the transformation system in *P. chrysosporium.* **A)** Schematic representation of the modifications in the GFP expression cassette (top) and the AMA1 autonomously replicating sequence (bottom). The upper panel shows the *GAPDH*’s exon1, intron1, and exon2 (eie) insertion and the four-nucleotide sequence ATGG (gray box) insertions between the *actin1* promoter and GFP coding region. Restriction sites utilized during the cassette construction and cloning are indicated. The lower panel shows the full-length AMA1 sequence (fAMA1) and the half-length AMA1 (hAMA1), which contains a single inverted repeat sequence and the unique central spacer sequence (UCS). **B)** Bright field and fluorescent micrographs of mycelium transformed with the six modified vectors. Scale bar: 100 µm. **C)** Comparison of GFP- associated fluorescence intensity in crude intracellular proteins driven by the modified *actin1* promoter regions in the six vector constructs. Mean ± SD values from three biological replicates are presented. **D)** The effect of vector design on transformant numbers. The number of transformants was estimated from the total colony count per 10^7^ protoplasts, adjusted based on the proportion of PCR-confirmed positives among 16 randomly selected colonies. Transformations were performed with 10 µg of each GFP expression vector, and protoplasts were cultured on CD agar media containing 0.3 µg/ml PT.

Transformation was performed using the newly constructed GFP expression vectors, and transformation efficiency, GFP expression in hyphae, and intracellular GFP levels were quantified (Fig. 3B-D), as described above. Each vector construct drove GFP expression in the mycelia, with fluorescence observed throughout the hyphal cells but not in the vacuoles, as visualized by fluorescence microscopy (Fig. 3B, Supplementary Fig. S3). Consistently, quantified intracellular fluorescence intensity levels from the new constructs were comparable with the original pPTRII- fAMA1-*actin1*pro-*egfp* vector (Fig. 3C). The inclusion of the ATGG sequence or the *eie* fragment, did not enhance gene expression in either the fAMA1 or hAMA1 constructs under the tested conditions (Fig. 3C). On the other hand, shortening the AMA1 sequence significantly improved the number of positive transformants (Fig. 3D). Notably, transformation efficiency of the original vector decreased from 28 to 14 (Fig 2D and 3D) compared to the previous experiment, which may be attributed to variability in protoplast quality, or experimental factors. To minimize this variability, the same batch of protoplasts was used for evaluating the impact of the improved constructs. Under these conditions, the improvement in transformation success was consistent for all hAMA1 constructs, increasing PCR positivity rates from 4-6 positive transformants (around 27%) to 12-14 positive transformants (around 81%) out of 16 randomly selected colonies (Fig. 3D, Supplementary Fig. S2). Respectively, we observed approximately 2-4 times increase in number of transformants for hAMA1 constructs.

## DISCUSSION

White-rot fungi are among the few organisms that excel in degrading recalcitrant lignin-rich wood materials. *P. chrysosporium* has long served as the standard white-rot basidiomycete for biochemical studies on lignin and polysaccharide degradation and has been heavily studied through proteomic and transcriptomic approaches (5, 32). Although genetic tools that enabling the direct investigation of gene function are becoming increasingly common in wood-decay fungi in recent years (26, 33–36), only a few early reports exist for *P. chrysosporium,* especially those that use dominant selectable markers, which are limited to transformation and homologous expression (14–16). To facilitate the functional analysis of genes involved in the decomposition of wood constituents or synthetic aromatic compounds in this fungus (15, 16), an optimized transformation protocol is a prerequisite. In the present study, we report a new transformation protocol for *P. chrysosporium,* utilizing *ptrA* as a dominant selection marker along with expression vectors constructed with native promoter elements.

In ascomycetes, pPTRII-based vector systems are well-established for genetic transformation (28) and have been successfully employed for heterologous gene expression (37) and genetic engineering in various *Aspergillus* species (21). However, there have been no reports of using either of these vectors or PT selection in basidiomycetes. Here, we demonstrated that *P. chrysosporium* growth is inhibited by PT and that the *A. oryzae*-derived resistance gene promotes colony growth (Fig. 1A, B). Selection with PT yielded a 100% PCR positivity rate in the resistant colonies for the pPTRII vector under optimal conditions, enabling a robust selection (Fig. 1C). Although the number of transformants obtained was relatively lower than that of *A. oryzae* (16.7 per μg of DNA per 10^7^ protoplasts) (28), our adapted protocol still yielded a sufficient number of transformants under optimized conditions (9.6 per μg of DNA per 10^7^ protoplasts, as shown in Table 1). This suggests that this method is applicable for *P. chrysosporium*, whilst the decrease in transformation success may be attributed to features inherent to the pPTRII plasmid sequence or differences in DNA uptake between these two phyla.

**Table 1.**
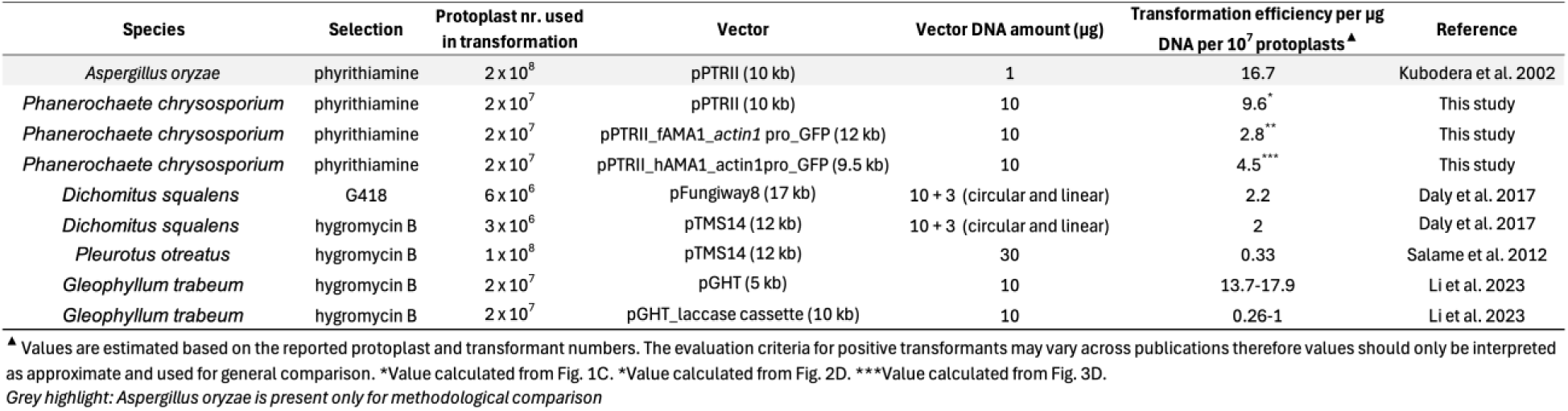
Comparison of transformation efficiency between wood-decaying basidiomycetes.

Heterologous expression is largely influenced by promoter activity in basidiomycetes (26, 38). Thus, we assessed the activity of four endogenous promoters - *actin1*, *DED*, *GAPDH*(1140), and *GAPDH*(1740) - using the *egfp* gene as a reporter. All promoters drove GFP expression detectable by fluorescent microscopy, out of which the *actin1* promoter displayed the strongest fluorescent signal and was selected for subsequent optimization (Fig. 2C). Notably, when the GFP cassette was inserted into pPTRII, the transformation efficiency decreased from 9.6 to 1.1-2.8, depending on the promoter sequence, which suggests that the insertion of the GFP expression cassette negatively affected transformation efficiency, probably due to the vector size (Table 1). The vector size dependency of transformation efficiency has been identified in other wood decaying basidiomycete *Gloeophyllum trabeum,* where doubling the vector size from 5 to 10 kb reduced the transformation efficiency to approximately one-fourth (33) (Table 1).

The insertion of introns in previous reports improved or was even indispensable for heterologous gene expression in some basidiomycete fungi (38–40). In particular, inserting a *GAPDH*-derived exon–intron–exon sequence at the start of the *egfp* gene, driven by the *GAPDH* promoter, led to a significant increase in GFP expression in *P. chrysosporium* (25). However, the same exon-intron-exon region inserted downstream of the *actin1* promoter had no considerable effect on GFP expression levels in our reaction setup (Fig. 3A-C). In addition, we speculated that inserting an 8 bp SbfI restriction site between the promoter and GFP start codon might affect gene expression by disrupting the native Kozak context at the 3′ end of the *actin1* promoter. Therefore, we fused a short ATGG sequence downstream of the *actin1* promoter, but despite this, no significant change was observed in GFP expression (Fig. 3A, C).

On the other hand, reducing the AMA1 sequence in the vector backbone allowed us to address the reduction in transformation efficiency associated with the increase in vector size. AMA1 functions as an autonomously replicating sequence in filamentous fungi, and its presence has been confirmed to improve transformation efficiency by 10-100 times in *A. niger* (41). That said, AMA1 is a considerably large sequence at 5.2 kb, making up more than half of the pPTRII vector. There have been efforts to shorten AMA1 for the ease of handling and to decrease vector load, which roughly halved the transformation efficiency in *A. niger* (42). However, we found that the number of transformants increased by an average of 3 times when one of the inverted repeats of the AMA1 sequence was deleted (hAMA), compared to the constructs that contained the full-length AMA1 (fAMA1) (Fig. 3A, D). The transformation efficiency of the pPTRII-hAMA1- *actin1*pro-*egfp* construct (4.5 transformants per μg of DNA per 10^7^ protoplasts) exceeded recent reports of PEG-mediated genetic transformation methods using conventional antibiotic-based selection methods for other wood-decaying basidiomycetes (Table 1). Transforming the white-rot fungus *Dichomitus squalens* using plasmids with G418 and hygromycin resistance markers yielded transformation efficiencies of 2.2 and 2, respectively (36). Using the same plasmid, the transformation efficiency in the white-rot *Pleurotus ostreatus* was reported to be approximately 0.22 (35). In the brown-rot basidiomycete, *G. trabeum,* transformation with a vector harboring the hygromycin resistance gene showed comparable transformation tendencies to our results, and transformation success was similarly influenced by the insertion of a laccase expression cassette (33). Although the integration status of the plasmid has not been explicitly tested, considering that the selection with PT falls within the range of methods conventionally used in basidiomycetes, it can serve as an effective transformation strategy for *P. chrysosporium*, and potentially for other basidiomycetes with high antibiotic tolerance, without relying on auxotrophic mutants. Additionally, pPTRII constructs have been successfully utilized for CRISPR-based genome editing in ascomycetes (19, 21), indicating the vector’s potential application beyond genetic transformation in *P. chrysosporium*.

Overall, the transformation protocol presented here offers a practical alternative to antibiotic-based selection, providing a tool for investigating lignocellulose-degrading pathways or other biotechnologically relevant molecular pathways in *P. chrysosporium*.

## MATERIALS AND METHODS

### Construction of GFP expression cassettes with native promoters

The pPTRII vector [Takara Bio Inc., Shiga, JAPAN], which harbors the *ptrA,* was used as a base for recombinant vector construction. For the promoter assays, GFP cassettes with four different types of promoters were constructed and inserted into the pPTRII vector. The GFP protein expressing gene, *egfp* (717 bp), was codon-optimized for expression in basidiomycetes [GenScript, Piscataway, NJ, USA]. The terminator region was constructed from the 3’ UTR region (194 bp) of the *P. chrysosporium glyceraldehyde-3-phosphate-dehydrogenase* (*GAPDH*) gene (JGI ProteinID: 6383341). Likewise, for promoter regions, native sequences upstream of the *actin1* gene (JGI Protein ID: 6238998), the *GAPDH* gene, and the *DED* gene (JGI ProteinID: 6206514) were selected. For the *actin1* promoter, we identified a 1037 bp region upstream of the *actin1* start codon, including the TATA box, to promote transcription initiation. Two variations of the *GAPDH* promoter were designed: a shorter version extending 1140 bp upstream of the *GAPDH* start codon and a longer version extending 1740 bp upstream, which includes the TATA box. As the final candidate, a 1715 bp region upstream of the *DED* gene was selected. The above four promoter regions were amplified from *P. chrysosporium* genomic DNA library by PCR using KOD-Plus-Neo polymerase [TOYOBO, Tokyo, Japan]. For genomic library preparation, mycelia were flash frozen in liquid nitrogen and crushed with a homogenizer [μT-12; Taitec Corporation, Saitama, Japan] at 2000 rpm for 15 seconds, repeated for a total of three cycles. The genomic DNA was extracted from the mycelial powder using the DNeasy Plant Mini Kit [QIAGEN, Venlo, the Netherlands]. KpnI and SbfI restriction sites were introduced via primer overhangs at the 5′ and 3′ ends of the amplified promoter regions, respectively (Supplementary Table S2). The promoters were cloned into pCR®Blunt II-TOPO® vector [Thermo Fisher Scientific Inc., Waltham, MA, USA]. The *egfp*-*GAPDH* 3’ UTR fragment, containing both the *egfp* gene and the terminator region, was DNA synthesized [GenScript, Piscataway, NJ, USA] and obtained in a pUC57 vector. The fragment was modified to contain a 5’ SbfⅠ site, a 3’ Hind III restriction site, and a NotⅠ restriction site separating the *egfp* gene from the terminator region. Subsequently, the four promoter regions were subcloned upstream of the GFP-*GAPDH* 3’ UTR fragment in the pUC57 vector using the KpnI and SbfI restriction sites to form the GFP expression cassettes. Then, GFP expression cassettes were inserted into the pPTRII vector using KpnI and HindIII, resulting in the following GFP expression vectors: pPTRII-*actin1*pro-*egfp,* pPTRII-*GAPDH*pro(1140)-*egfp*, pPTRII- *GAPDH*pro(1740)-*egfp*, and pPTRII-*DED*pro-*egfp*.

### Modifying the GFP expression vector

In an attempt to enhance GFP expression, two types of short DNA elements were inserted between the promoter region and the GFP coding sequence of the pPTRII-*actin1*pro-*egfp* vector.

Either a four-nucleotide sequence (ATGG) or an eie sequence consisting of the exon 1 (6 bp), intron 1 (55 bp), and exon 2 (9 bp) of the GAPDH gene: ATGCCGgtcagtacaccacacagcccgaccgcgacgaccgcgtgctgacttcgctttccagGTCAAAGCA (Fig. 3A, top) (25). These sequences were inserted into the vector by PCR amplification using reverse primers containing the inserts and an SbfI restriction enzyme site (Supplementary Table S2) (Fig. 3A).

To improve the transformation efficiency, the full-length AMA1 sequence (fAMA1) within the pPTRII vector was shortened to approximately half of its original length (hAMA1) by using inverse PCR (Supplementary Table S2) (Fig. 3A). The following vectors were produced: pPTRII-hAMA1-*actin1*pro-*egfp*, pPTRII-fAMA1-*actin1*pro-eie-*egfp*, pPTRII-hAMA1- *actin1*pro-eie-*egfp*, pPTRII-fAMA1-*actin1*pro-ATGG-*egfp*, pPTRII-hAMA1-*actin1*pro-ATGG- *egfp*.

### Protoplast isolation

*P. chrysosporium* RP-78 strain was used for this study. Protoplasts were prepared using a slightly modified version of the protocol described by Stewart et al. (43). 1 × 10⁷ spores of *P. chrysosporium* were inoculated into 100 mL of YMPG liquid medium and statically cultured at 37°C for 40 hours. Mycelia were collected and cell walls were digested with 10 mg/mL Yatalase [Takara] and Cellulase Onozuka R-10 [Wako] in 10 mM sodium phosphate buffer (pH 6.0) containing 0.8 M NaCl for 3 hours at 37℃ with shaking at 120 rpm. The collected protoplasts were gently resuspended and adjusted to 1 × 10⁸ protoplast/mL in Solution 1 (0.8 M NaCl and 10 mM CaCl_2_ in 10 mM Tris-HCl [pH 8.0]), then additional 0.2 volumes of Solution 2 (40% [w / v] PEG4000 and 50 mM CaCl_2_ in 50 mM Tris-HCl [pH 8.0]) was added.

### Growth inhibition of *P. chrysosporium* by PT

Approximately 2 x 10^6^ protoplasts of *P. chrysosporium* wild type strain were cultured at 26.5°C for 4 days on CD agar plates containing PT of various concentrations (0 ∼ 1.0 μg/mL), and the number of recovered colonies was counted (Supplementary Table S1).

### Fungal transformation

The transformation method used in this study basically follows the manufacturer’s instructions of the pPTRII vector [Takara]. To determine the optimal transformation conditions, 1, 5, or 10 μg of donor DNA vector was added to 200 μL protoplast suspension (approximately 2 × 10^7^ protoplasts) and incubated on ice for 30 minutes. An additional 1 mL of Solution 2 was added, gently suspended, and incubated for 15 minutes at room temperature. Transformed protoplasts were diluted by 8.5 mL of Solution 1, gently mixed, and collected by centrifugation for 10 minutes at 2000 rpm. Pelleted protoplasts were resuspended in regeneration media (100 μL of Solution 1 supplemented with 1% glucose), and incubated for 1 hour at 37 °C with shaking at 120 rpm. For selection, protoplasts were plated on CD agar medium containing 0.1, 0.3, 0.5, or 1.0 μg/mL PT and cultured at 26.5 °C for 3-4 days.

### Genomic PCR

Genomic DNA was extracted from the mycelia of PT-resistant colonies, exhibiting growth in the presence of appropriate PT concentrations. To confirm the proper introduction of the pPTRII vector and its GFP constructs, PCR amplification was performed using KOD FX Neo polymerase [TOYOBO]. For the empty pPTRII, a vector backbone-specific primer set was used, and for the GFP constructs, an *egfp*-specific primer set was used. The nuclear ribosomal ITS region was amplified as a reference for genomic DNA loading amount with specific primers (for primer sequences see Supplementary Table S2). Amplicons were analyzed by 1% agarose gel electrophoresis.

### Calculation of the number of transformants and transformation efficiency

To estimate the *number of transformants* per 10⁷ protoplast, the total number of PT-resistant colonies was counted and corrected based on the PCR positivity rate. The PCR positivity rate was defined as the proportion of genomic PCR positives among the total number of screened colonies. Transformation efficiency was calculated as the *number of transformants* per µg of donor DNA, using the following formula: Transformation efficiency = *number of transformants* / µg donor DNA.

### Observation of transformed mycelium with fluorescence microscopy

GFP-expressing transformants were grown on CD agar plates with 0.3 μg/mL PT, and the GFP fluorescence of the hyphae was observed and photographed by a fluorescence microscope [DM2500, Leica] equipped with the camera [DFC310 FX, Leica]. L5 filter cube was used for GFP detection (excitation filter: BP 480/40; dichromatic mirror: 505; suppression filter: BP 527/30 nm).

### Determination of intracellular GFP expression

For determining intracellular GFP expression, a previously reported protocol (25) was slightly modified. For liquid cultures, ten agar plugs of each mycelia transformed with either GFP expression vectors or the pPTRⅡ vector were inoculated into 50 mL of CD liquid medium containing 0.3 μg/mL PT and incubated at 37°C for 5 days. Mycelia were harvested, frozen in liquid nitrogen, homogenized with a crusher, and suspended in TE + 0.002 % NaN3 buffer. Following centrifugation, the supernatants of the crude intracellular proteins were collected, and total protein contents were assayed by the Bradford method [Protein Assay, Bio-Rad, Hercules, CA, USA]. The protein concentration was adjusted to 1 mg/mL with TE + 0.002 % NaN3 buffer. The fluorescent intensity of the intracellular protein extract was measured using a fluorescence spectrophotometer [F-2500, Hitachi, Tokyo, Japan] with an excitation wavelength of 488 nm and an emission wavelength of 509 nm.

## Acknowledgments

This work was supported by JST FOREST program (JPMJFR233T), JSPS KAKENHI (JP22K19195 and JP23K21222), Sugiyama Chemical & Industrial Laboratory, Institution of Fermentation Osaka, Noda Institute for Scientific Research, and The Naito Foundation to CH.

## Author contributions

K.M. carried out the experiments, analyzed the data, and wrote the draft. B.P. analyzed the data and wrote the draft. A.Y. carried out vector constructions. M.K. wrote the draft. C.H. designed the research, carried out experiments, analyzed the data and wrote the draft. All authors reviewed and approved the paper. There is no conflict of interest.

